# In-gel protein digestion using acidic methanol produces a highly selective methylation of glutamic acid residues

**DOI:** 10.1101/2024.03.15.585290

**Authors:** Marta Lozano-Prieto, Emilio Camafeita, Inmaculada Jorge, Andrea Laguillo-Gómez, Rafael Barrero-Rodríguez, Cristina A. Devesa, Clara Pertusa, Enrique Calvo, Francisco Sánchez-Madrid, Jesús Vázquez, Noa B. Martin-Cofreces

**Affiliations:** Immunology Service, Hospital Universitario de la Princesa, Universidad Autónoma de Madrid (UAM), Instituto Investigación Sanitaria Princesa (IIS-IP), Madrid, Spain; Cardiovascular Proteomics Laboratory, Centro Nacional de Investigaciones Cardiovasculares Carlos III (CNIC), Madrid, Spain; Centro de Investigación Biomédica en Red, Enfermedades Cardiovasculares (CIBERCV), Madrid, Spain; Intercellular Communication in the Inflammatory Response. Vascular Pathophysiology Area. Centro Nacional de Investigaciones Cardiovasculares (CNIC), Madrid, Spain; Universidad Autónoma de Madrid, Madrid, Spain; Videomicroscopy Unit, Instituto de Investigación Sanitaria La Princesa (IIs-Princes), Madrid, Spain

## Abstract

Mass-tolerant open search methods allow the high-throughput analysis of modified peptides by mass spectrometry. These techniques have paved the way to unbiased analysis of post-translational modifications (PTMs) in biological contexts, as well as of chemical modifications produced during the manipulation of protein samples. In this work, we have analyzed in-depth a wide variety of samples of different biological origin, including cells, extracellular vesicles, secretomes, centrosomes and tissue preparations, using Comet-ReCom, a recently improved version of the open search engine Comet-PTM. Our results demonstrate that glutamic acid residues undergo intensive methyl esterification when protein digestion is performed using in-gel techniques, but not using gel-free approaches. This effect was highly specific to Glu and was not found for other methylable residues such as Asp.

---

PTMs are known to exert a pivotal role in the control of signaling cascades and protein structure and functionality by regulating protein localization, activity, folding, interaction and stability, and in recent years have been implicated in pathological processes as tumorigenesis [1], neurodegeneration [2, 3] and atherosclerosis [4]. While mass spectrometry (MS) is the method of choice for PTM identification and quantitation, conventional (also termed *closed*) database search approaches followed in proteomics research rely on matching exact mass differences, therefore restricting PTM analyses to a few modifications (typically less than four) to keep a reasonably sized search space. This limitation has been circumvented in recent years by mass-tolerant database searching methods [5] such as MSFragger [6], Comet-PTM [7] or Comet-ReCom, a recently improved version of the former [8].

Re-analyses with Comet-ReCom of the LC-MS/MS data from centrosomal-enriched fractions obtained from human T lymphocytes [9] to study the presence of PTMs revealed a strikingly high proportion of the total peptide-spectrum matches (PSMs) bearing a +14.01565 Da modification (corresponding to methylation, (see Suppl. Fig. 1) at Glu residues (3,48% with this modification) (see ‘proteome identifier 5’ in Fig. 1B). These samples were processed using the in-gel digestion procedure, a technique that is advantageous in some contexts because it uses the gel matrix as a reaction chamber to eliminate contaminants or for the addition and washing of reagents [10]. There is previous preliminary evidence, based on MS analysis of a very limited number of peptides (5) that the acidified methanol solutions used for gel staining/destaining may produce artifactual methylation of Glu and, to a lesser extent, of other residues such as Asp [11]. This is a well-known reaction whereby a carboxylic acid undergoes esterification by an alcohol in acidic medium. However, in our case the methylation appeared to be highly specific to Glu, being virtually undetectable in Asp (<0.1%, 35-fold lower than Glu) or other residues amenable to methylation (*i*.*e*. Lys, Arg, Glu, His, Asn, Gln, and Cys) (‘proteome identifier 5’ in Fig. 1B). This was particularly remarkable given the high similarity between Glu and Asp, which differ only in the presence of an extra methylene group in the side chain of the former. We therefore decided to take advantage of open search methods to analyze this effect in more detail in a wide range of samples of different origins.

**Figure 1:**
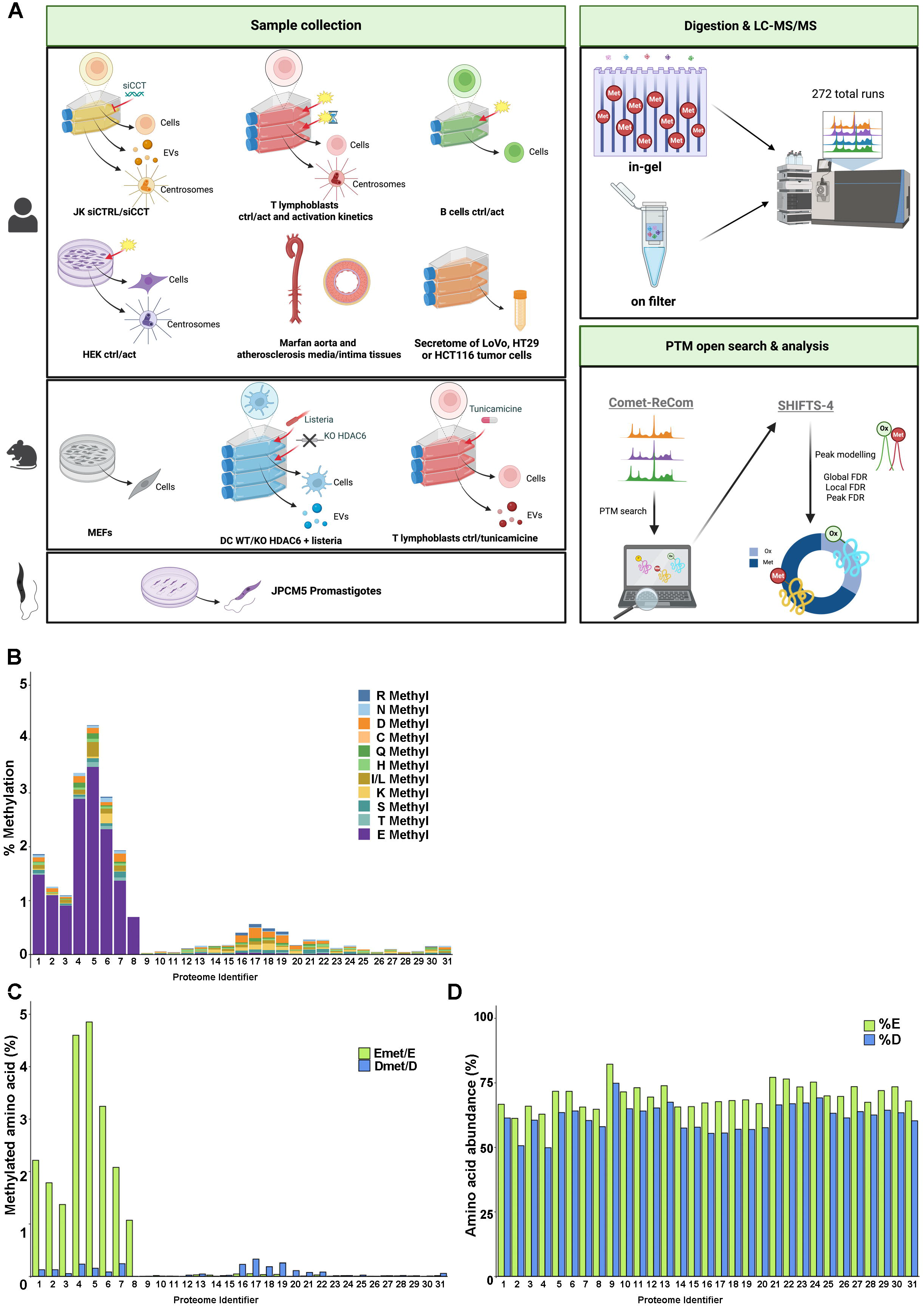
**(A)** Schematic representation of the unbiased PTM analysis based on Comet-ReCom open database search, performed with different cell, extracellular vesicle (EV), secretome, centrosome and tissue preparations digested in-gel or on-filter. **(B)** Amino acid distribution in methylable residues of the +14.01565 Da modification across the cell, extracellular vesicle, secretome, centrosome and tissue samples digested in-gel or on-filter (see Supplementary Information). **(C)** Percentage Glu and Asp residues bearing the +14.01565 Da modification across the cell, extracellular vesicle, secretome, centrosome and tissue samples digested in-gel or on-filter. **(D)** Percentage of PSMs containing Asp or Glu across samples.

We applied Comet-ReCom to the analysis of distinct cell, extracellular vesicle, secretome, centrosome and tissue preparations digested either in-gel (using acidified methanol in the staining/destaining procedure) [12] or on-filter (without methanol) [13] (Fig. 1A, see Supplementary Table 1). We found a considerably higher prevalence of the +14.01565 Da modification at Glu residues in the proteomes digested in-gel as compared to on-filter digestion (Fig. 1B), but not at Asp or other methylable residues, strongly suggesting that the modification produced by in-gel treatments was highly specific for Glu. Of note, the prevalence of Asp residues bearing the +14.01565 Da modification was very low in comparison and was not affected by the digestion method (Fig. 1C). Note also that the relative abundance of Glu and Asp residues was quite similar across all the proteomes analyzed (Fig. 1D), ruling out the possibility that this effect was due to a higher Glu abundance in these samples. To further verify the Glu specificity towards the in-gel digestion artefact, we applied Comet-Recom to LC-MS/MS data obtained from the same set of mouse embryonic fibroblast samples digested using both in-gel and on-filter protocols [14]. The results confirmed that the in-gel protocol produces a highly specific methyl esterification of Glu but not of other methylable residues (see ‘proteome identifiers 6 and 30’ in Fig. 1B).

The markedly different reactivity of Glu and Asp residues towards methanol in acidic media may be attributed to the distinct interaction of the side-chain carboxy group with the peptide backbone [11]. It is noteworthy that, despite their chemical similarity, significant differences between Glu and Asp reactivity have been recently reported using molecular dynamics simulations [15]. Regarding the other methylable residues, the reaction of methanol with His, Lys and Arg residues is hindered in acidic media due to the protonation of their side chains. These results were also observed in other proteomic data generated by other research groups (Fig. 1B protein identifiers 7, 8 and 31) (16,17). Our observation that in-gel digestion methods produce a highly specific methylation of Glu will serve the proteomics community to better interpret open search-based PTM analysis performed using this technique.

## Acknowledgments

Fig. 1A was created with Biorender. This study was supported by the Spanish Ministry of Science and Innovation, Agencia Estatal de Investigación by competitive grants PID2021-122348NB-I00, PID2022-141890B-I00, PID2020-120412RB-I00, PDC2021-121797-I00 and PGC2018-097019-BI00 funded by MICIU/AEI/ 10.13039/501100011033 and by “ERDF A way of making Europe”, PLEC2022-009298, PLEC2022-009235 and EQC2021-007053-P funded by MICIU/AEI/10.13039/501100011033 and by “European Union NextGenerationEU/ PRTR”, and S2022/BMD-7333-CM (INMUNOVAR-CM) and P2022/BMD7209 (INTEGRAMUNE) funded by Comunidad de Madrid. CIBER Cardiovascular (CB16/11/00272, CB16/11/00277) Fondo de Investigación Sanitaria del Instituto de Salud Carlos III; co-funding by Fondo Europeo de Desarrollo Regional (FEDER). The project leading to these results has received funding from “La Caixa” Foundation under the project codes LCF/PR/HR22/52420019 and LCF/PR/HR23/52430018. MLP is supported by a FPI fellowship (PRE2021-097478). ALG is supported by a FPU fellowship (FPU18/03882). RBR is supported by a FPU fellowship (FPU20/03365). CAD is supported by a FPI fellowship (PRE2019-090019). The CNIC is supported by the Instituto de Salud Carlos III (ISCIII), the Ministerio de Ciencia, Innovación Y Universidades (MICIU) and the Pro CNIC Foundation), and is a Severo Ochoa Center of Excellence (grant CEX2020-001041-S funded by MICIU/AEI/10.13039/501100011033).

## Author contribution

MLP, data curation, formal analysis, validation, writing of original draft, Fig 1; EC, data curation, formal analysis, validation, writing of original draft; IJ, data curation, formal analysis, validation, writing of original draft; ALG, software; RBR, software; CAD, software; CP, data curation; EC, data curation, formal analysis; FSM, JV and NBMC: conceptualization, funding, supervision, writing of original draft.

## Data availability

All data included in this study will be available (see Suppl. Material).

## References

1. Liu, R., et al., mTORC1 activity regulates post-translational modifications of glycine decarboxylase to modulate glycine metabolism and tumorigenesis. Nat Commun, 2021. 12(1): p. 4227.

2. Alquezar, C., S. Arya, and A.W. Kao, Tau Post-translational Modifications: Dynamic Transformers of Tau Function, Degradation, and Aggregation. Front Neurol, 2020. 11: p. 595532.

3. Junqueira, S.C., et al., Post-translational modifications of Parkinson’s disease-related proteins: Phosphorylation, SUMOylation and Ubiquitination. Biochim Biophys Acta Mol Basis Dis, 2019. 1865(8): p. 2001–2007.

4. Pirillo, A., et al., Impact of protein glycosylation on lipoprotein metabolism and atherosclerosis. Cardiovasc Res, 2021. 117(4): p. 1033–1045.

5. Chick, J.M., et al., A mass-tolerant database search identifies a large proportion of unassigned spectra in shotgun proteomics as modified peptides. Nat Biotechnol, 2015. 33(7): p. 743–9.

6. Yu, F., et al., Identification of modified peptides using localization-aware open search. Nat Commun, 2020. 11(1): p. 4065.

7. Bagwan, N., et al., Comprehensive Quantification of the Modified Proteome Reveals Oxidative Heart Damage in Mitochondrial Heteroplasmy. Cell Rep, 2018. 23(12): p. 3685–3697 e4.

8. Laguillo-Gomez, A., et al., ReCom: A semi-supervised approach to ultra-tolerant database search for improved identification of modified peptides. J Proteomics, 2023. 287: p. 104968.

9. Martin-Cofreces, N.B., et al., The chaperonin CCT controls T cell receptor-driven 3D configuration of centrioles. Sci Adv, 2020. 6(49).

10. Martinez-Acedo, P., et al., A novel strategy for global analysis of the dynamic thiol redox proteome. Mol Cell Proteomics, 2012. 11(9): p. 800–13.

11. Haebel, S., et al., Electrophoresis-related protein modification: alkylation of carboxy residues revealed by mass spectrometry. Electrophoresis, 1998. 19(5): p. 679–86.

12. Bonzon-Kulichenko, E., et al., A robust method for quantitative high-throughput analysis of proteomes by 18O labeling. Mol Cell Proteomics, 2011. 10(1): p. M110 003335.

13. Wisniewski, J.R., et al., Universal sample preparation method for proteome analysis. Nat Methods, 2009. 6(5): p. 359–62.

14. Bonzon-Kulichenko, E., et al., Improved integrative analysis of the thiol redox proteome using filter-aided sample preparation. J Proteomics, 2020. 214: p. 103624.

15. Lemke, T., et al., Three Reasons Why Aspartic Acid and Glutamic Acid Sequences Have a Surprisingly Different Influence on Mineralization. J Phys Chem B, 2021. 125(36): p. 10335–10343.

16. Almeida-Marques, C., et al., Secretome processing for proteomics: A methods comparison. Proteomics, 2024: p. e2300262.

17. Sanchiz, A., et al., The Experimental Proteome of Leishmania infantum Promastigote and Its Usefulness for Improving Gene Annotations. Genes (Basel), 2020. 11(9).

