## Supplementary figures and images for "In-gel protein digestion using acidic methanol produces a highly selective methylation of glutamic acid residues"

### Supplemental Fig 1

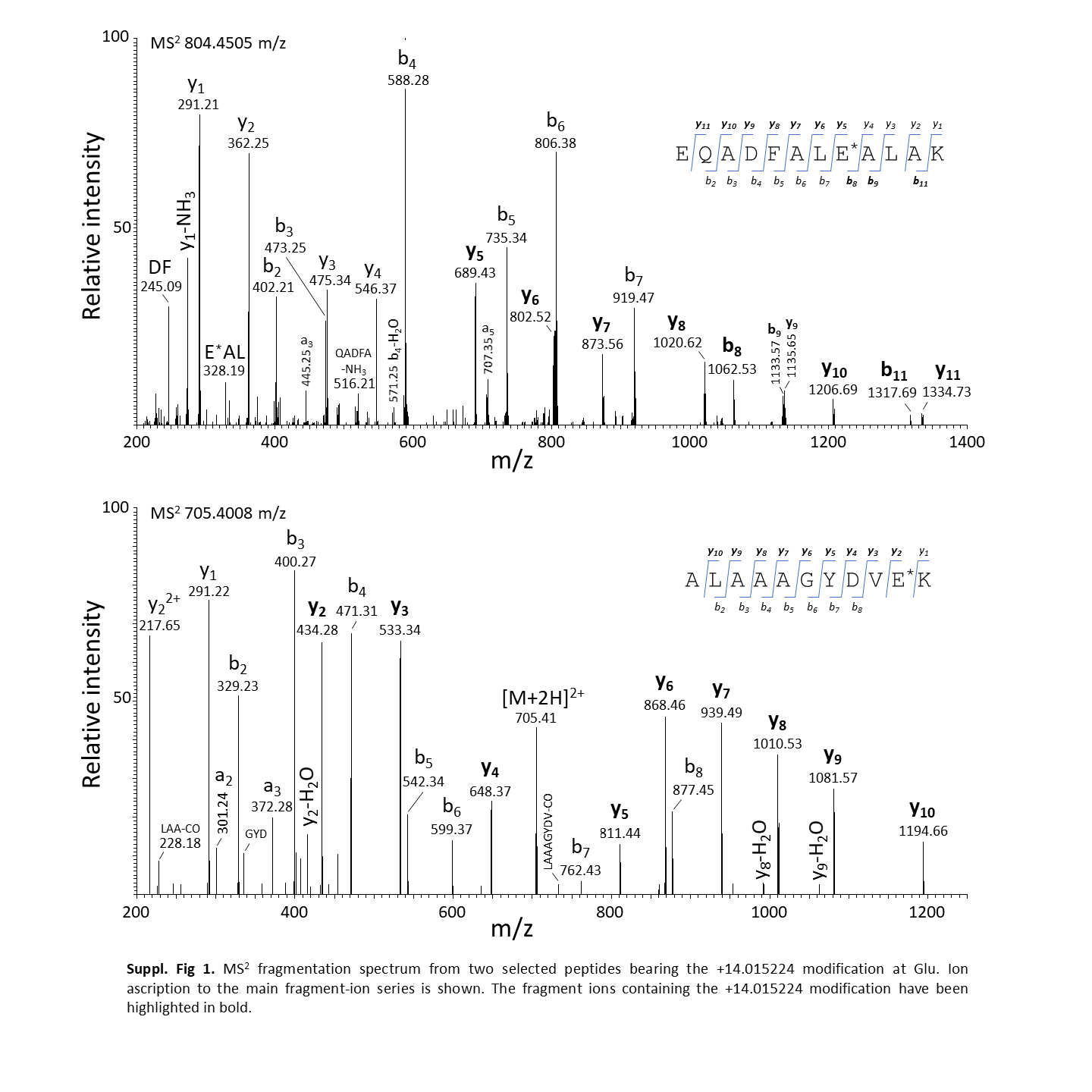
