## Supplemental Methods for "In-gel protein digestion using acidic methanol produces a highly selective methylation of glutamic acid residues"

**SUPPLEMENTARY METHODS**

1. **MS DATASETS**

Raw LC-MS/MS data from a total of 192 samples comprising the following cell, extracellular vesicle/exosome, secretome, centrosome and tissue preparations were used in this work:

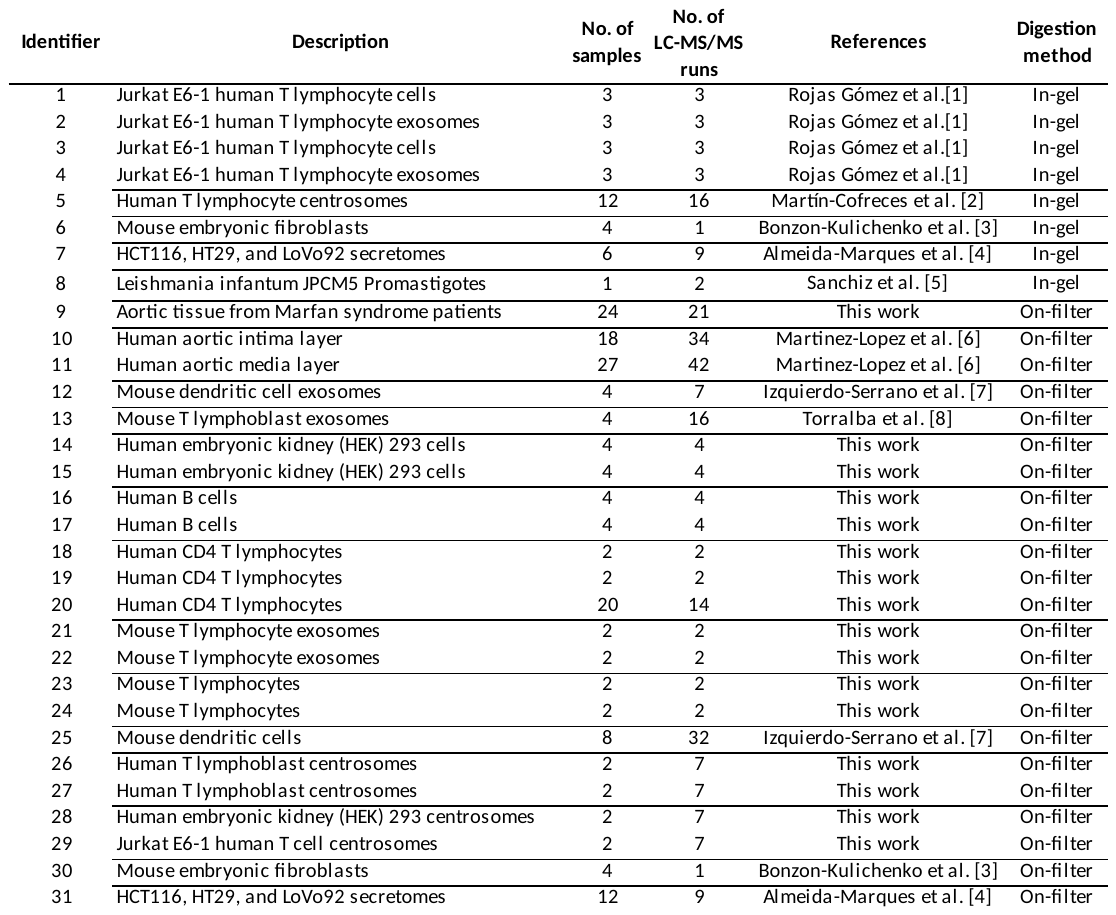

1.1. **MS data retrieved from previously published work**

LC-MS/MS data from previously published work using in-gel or on-filter digestion were used for the following samples:

1.1.1. **In-gel digested samples**

Jurkat E6-1 human T lymphocyte lysates and exosomes (samples 1-4) [1], human T lymphocyte centrosomes (sample 5) [2], mouse embryonic fibroblasts (sample 6) [3], HCT116, HT29, and LoVo92 secretomes (sample 7) [4], and *L. infantum* JPCM5 Promastigotes (sample 8) [5].

1.1.2. **On-filter digested samples**

Human aortic intima and media layers (samples 10 and 11) [6], mouse dendritic cell lysates and exosomes (samples 12 and 25) [7], mouse T lymphoblast exosomes (sample 13) [8], and mouse embryonic fibroblasts (sample 30) [3], and HCT116, HT29, and LoVo92 secretomes (sample 31) [4].

1.2. **MS data obtained in this work**
     1.2.1. **Ethics statement**

Animal works have been conducted according to approved procedures by the local Ethics Committee for Basic research at the CNIC Ethical Committee for Animal Welfare and the Organo Encargado del Bienestar Animal (OEBA) del Gabinete Veterinario de la Universidad Autónoma de Madrid (UAM). All of them are in agreement with EU Directive 86/609/EEC and Recommendation 2007/526/EC regarding the protection of animals used for experimental and other scientific purposes, enforced in Spanish law under Real Decreto 53/2013 (Authorization ProEX 206.1/20).

Human samples 16-20, 26 and 27 were isolated from buffy coat preparations from healthy donors provided by Centro de Transfusiones de la Comunidad de Madrid under an agreement with the IIS-Princesa (Madrid, Spain) in accordance with the Declaration of Helsinki and the government ethical consent, and approved by the Hospital La Princesa Research Ethics Committee.

Aortic tissue samples from Marfan syndrome patients (sample 9) were provided by the biobanks of Hospital Vall d'Hebron (Barcelona, Spain), Puerta de Hierro (Madrid, Spain) and Valdecilla (Santander, Spain) under authorization of the Instituto de Salud Carlos III Ethics Committee (CEI PI 91_2018-v2-Enm_2019_Amp 2023).

1.2.2. **Materials**
       1.2.2.1. **Mouse T lymphocyte cells and exosomes** (samples 21-24) were prepared as described in Torralba *et al*. [8].
       1.2.2.2. **Jurkat E6-1 human T cell centrosomes** (sample 29). Centrosome-enriched subcellular fractions from Jurkat E6-1 cell line were prepared as described in [2]. In brief, cells were ice-cooled, centrifuged at 2,000g at 25 °C for 30 min, washed in Tris Buffered Saline and resuspended in lysis buffer (2 × 10^6^ cells/ml) 1 mM tris-HCl (pH 7.4) containing 0.5% NP-40, 0.5 mM MgCl_2_, 0.1 % β-mercaptoethanol, and inhibitors of proteases and phosphatases. After 10 min with gentle shaking, lysates were filtered through a nylon mesh and centrifuged at 24,000 xg for 10 min at 4 °C. The supernatant fraction was filtered and re-equilibrated into 10 mM piperazine-N,N-bis(2-ethanesulfonic acid) (PIPES) (pH 7.2) and centrifuged at 24,000 xg for 60 min at 4 °C to pellet the centrosomes. Centrosome-enriched pellet was resuspended in gradient buffer (10 mM PIPES (pH 7.2) containing 5% sucrose, 0.1% β-mercaptoethanol and 0.1% Triton X-100) and homogenized through 21-gauge (four times) and 25-gauge (five times) needles before loading the 10 mL onto the first sucrose gradient, including (bottom-top) 1 mL 70%, 2 mL 50%, 2 mL 40%, 2 mL 30%, and 2 mL 20% (w/w). This gradient was subjected to a 32,500 xg centrifugation for 1 h at 4°C (SW28 rotor). Collection of fractions was performed from the top and 50 to 70% were collected together and diluted with gradient buffer to a final concentration of about (20-30 %) of sucrose to load the sample onto a second sucrose gradient including (bottom-top) 0.5 mL 70%, 0.5 mL 50%, and 1 mL 40% (w/w) that was centrifuged at 130,000 xg for 1 hour at 4°C (SW40 Ti rotor). Fractions of 200 μL were recovered from top and analyzed for γ-tubulin content by Western blot and for quantitation of total protein through colorimetric detection by BCA Protein Assay.
       1.2.2.3. **Human embryonic kidney (HEK) 293 cells and centrosomes** (samples 14, 15 and 28). HEK 293 cells were scrapped from cell culture plates and 10^9^ cells were processed as described above to isolate centrosome-enriched fractions.
       1.2.2.4. **Human B cells** (samples 16 and 17). Primary human B lymphocytes were isolated from Buffy coats through a ficoll gradient to obtain peripheral blood mononuclear cells and later through negative selection with magnetic beads (human B Cell negative kit from StemCell Technologies). Cell were stimulated or not with IL-4 and LPS for 4 h, centrifuged at 5,500 rpm for 5 min at 4 °C and dry pellets were stored at -80 °C until processed to obtain protein extracts.
       1.2.2.5. **Human CD4 T lymphocytes** (samples 18-20). Primary human T CD4 lymphocytes were isolated from Buffy coats through a ficoll gradient to obtain peripheral blood mononuclear cells and later through negative selection with magnetic beads (human CD4 T Cell negative kit from StemCell Technologies). Cells were then stimulated with Immunocult Activator CD3CD28 (STemCell Technologies)or not for the times indicated, centrifuged at 5,500 rpm for 5 min at 4°C and dry pellets were stored at -80 °C until processed to obtain protein extracts.
       1.2.2.6. **Human T lymphoblast centrosomes** (samples 26 and 27). Primary human T lymphoblasts were generated by stimulating peripheral blood lymphocytes (PBLs) isolated from Buffy coats through a ficoll gradient and an adhesion step to deplete monocytes. PBLs were cultured in RPMI supplemented with 10 % FCS and activated with Staphylococcus enterotoxin E (SEE; 0.01 µg/mL) and phytohaemagglutinin (PHA; 0.2 µg/mL) for 48 h. Cells were then washed with Hank’s balanced salt solution (HBSS) and grown in RPMI including 10 % FCS and IL-2 (25 U/mL) for 7-10 days. Cells from same donor were split in two and stimulated with monoclonal antibodies anti-CD3ε and anti-CD28-coated beads or gamma-globulins-coated beads as control for 15-20 min. Cells were then ice-cooled and subjected to the centrosome-enriched fraction isolation procedure as above. Samples from different donors were pooled after first lysis.
       1.2.2.7. **Aortic tissue from patients with Marfan syndrome (**sample 9). Control samples of ascending aorta were obtained from organ transplant donors, and aortic samples from Marfan syndrome patients were obtained in the course of elective or emergency aortic surgery.
1.2.3. **Preparation of protein extracts**

Whole cell, exosome and centrosome preparations (samples 14-24 and 26-29) were boiled at 95 °C for 5 min in lysis buffer (50 mM Tris-HCl pH 6.8, 2% SDS, 10 mM DTT). Human aortic tissue samples (sample 9) were homogenized using a FastPrep-24 instrument (MP Biomedicals, Illkirch, France), following the manufacturer’s instructions, and the resulting homogenates were boiled for 5 min at 95 °C . The samples were centrifuged at 12,000 rpm at 4^o^C for 15 min to remove debris and the supernatants were collected. Protein concentration in the supernatant fractions was measured with the RC/DC protein assay kit (BioRad) according to the manufacturer’s instructions.

1.2.4. **On-filter protein digestion**

Protein samples were mixed with denaturing buffer (8 M urea in 100 mM Tris-HCl pH 8.5) and concentrated on 30 K FASP filters (Expedeon). After washing with denaturing buffer at 10,000 rpm for 15 min, free thiol groups were alkylated by incubation with 50 mM iodoacetamide. Then the filters were washed with denaturing buffer followed by washing with trypsin digestion buffer (50 mM ammonium bicarbonate pH 8.8). Protein samples were digested overnight at 37°C with sequencing grade trypsin (Promega, Madison, WI, USA) at 1:40 (w/w) trypsin:protein ratio in digestion buffer, after which the resulting tryptic peptides were recovered by centrifugation. Trifluoroacetic acid was added to a final concentration of 1% and the peptides were desalted on C18 Oasis HLB extraction cartridges (Waters Corporation, Milford, MA, USA) and dried-down.

1.2.5. **Peptide isobaric labeling**

The dried peptides corresponding to a set of 20 human T lymphocyte cells (sample 20) and the 24 aortic tissue samples from patients with Marfan syndrome (sample 9) were taken up in 100 mM triethylammonium bicarbonate, and peptide concentration was determined using a Direct Detect IR spectrometer (Millipore, Billerica, MA, USA). Peptide samples were labeled with tandem mass tags (TMT, Thermo Fisher Scientific, Waltham, MA, USA) according to the manufacturer’s instructions. The labelled samples were distributed across two and three TMT 10-plex experiments for human T lymphocyte and aortic tissue samples, respectively, desalted using C18 Oasis HLB extraction cartridges (Waters) and dried-down.

1.2.6. **LC-MS/MS analysis**

The dried peptide samples were taken up in 0.1% formic acid for LC-MS/MS analysis.

1.2.6.1. **Mouse T lymphocyte cell and exosome samples** (samples 21-24) were applied to an EASY-nLC 1000 nano-flow HPLC system (Thermo Fisher Scientific) coupled on-line with a Q Exactive mass spectrometer (Thermo Fisher Scientific). C18-based reverse phase separation was used with a 2-cm trap column and a 50-cm analytical column (EASY-Spray, Thermo Fisher Scientific). Peptides were loaded in buffer A (0.1% formic acid (v/v)) and eluted with a 180-min linear gradient of buffer B (90% ACN, 0.1% formic acid (v/v)) at 200 nL/min flow. Mass spectra were acquired in a data-dependent manner, with an automatic switch between MS and MS/MS using a top 15 method and 45 s dynamic exclusion. The MS spectra were acquired with the orbitrap analyzer in the 400–1,500 m/z mass range with 70,000 resolution, and higher-energy collisional dissociation (HCD) peptide fragments obtained at 30 normalized collision energy were analyzed with 35,000 resolution in the orbitrap.
1.2.6.2. **Human HEK 293 cells, B cells and T lymphocytes** (samples 14-20) were applied to an EASY-nLC 1000 nano-flow HPLC system coupled on-line with a quadrupole orbitrap Q Exactive HF mass spectrometer (Thermo Fisher Scientific). C18-based reverse phase separation was used with a 2-cm trap column and a 50-cm analytical column (EASY-Spray). Peptides were loaded in buffer A (0.1% formic acid (v/v)) and eluted with a 180-min linear gradient of buffer B (90% AcN, 0.1% formic acid (v/v)) at 200 nL/min flow. Spectra were acquired using full ion-scan mode over the 400-1,500 m/z range and 120,000 resolution. The automatic gain control (AGC) target was set at 2 x 105 with a maximum injection time of 50 ms. Data-dependent speed mode acquisition was performed at 5 x 104 AGC and 120 ms injection time, with a 1-Da isolation window and 45 s dynamic exclusion. HCD fragmentation was set to 33% normalized collision energy. MS/MS scan resolution was set to 30,000.
1.2.6.3. **Human centrosome samples from T lymphoblasts and HEK 293 and Jurkat E6-1 cells** (samples 26-29) were analyzed on the Evosep One system (Evosep, Odense, Denmark) using the preprogrammed 88-min gradient essentially as described [9]. MS was performed with an Orbitrap Eclipse (Thermo Fisher Scientific) in positive ion mode using data-dependent acquisition with 2 s top speed cycles. Each cycle consisted of one full MS scan followed by as many MS/MS events that could fit within the given 2 s cycle time limit. MS scans were collected at a resolution of 120,000 (410–1600 m/z range, 4 x 105 AGC, 50 ms maximum ion injection time). HCD MS/MS spectra were acquired at a resolution of 30,000 (0.7 m/z isolation width, 35% collision energy, 1.25 x 105 AGC target, 54 ms maximum ion time). Dynamic exclusion was set to exclude previously sequenced peaks for 20 s within a 10-ppm isolation window.
1.2.6.4. **Aortic tissue samples from Marfan syndrome patients** (sample 9) were applied to an EASY-nLC 1000 nano-flow HPLC system coupled on-line with an orbitrap Fusion mass spectrometer (Thermo Fisher Scientific). C18-based reverse phase separation was used with a 2-cm trap column and a 50-cm analytical column (EASY-Spray). Peptides were loaded in buffer A (0.1% formic acid (v/v)) and eluted with a 300-min linear gradient of buffer B (90% ACN, 0.1% formic acid (v/v)) at 200 nL/min flow. Mass spectra were acquired in a data-dependent manner, with an automatic switch between MS and MS/MS, using a top-speed method and 30 s dynamic exclusion. MS spectra were acquired in the 400–1,500 m/z range at 120,000 resolution, while HCD MS/MS were performed at 33 normalized collision energy and analyzed with 35,000 resolution in the orbitrap.

2. **MS DATA ANALYSIS**

The Thermo Scientific LC-MS/MS RAW files retrieved from previous analyses or obtained in this work (272 total runs) were converted to the mzML format using ThermoRawFileParser version 1.1.9 [10]. The mzML MS data were open-searched against a mouse or human Uniprot (May 2021) concatenated target-decoy database with Comet-ReCom [11] using the following parameters: one maximum trypsin missed cleavage site, 500 Da precursor mass tolerance, 0.02 Da fragment mass tolerance. Cys carbamidomethylation was in all cases considered as a fixed modification. Additionally, in the case of the samples subjected to isobaric labeling (see Subsection 1.2.5 above), the corresponding mass increments were set as fixed modifications at Lys and peptide N-terminus: 144.102063 Da (iTRAQ 4-plex) for the in-gel digested human T lymphocyte centrosomes (sample 5) [2] and the mouse dendritic cell exosomes (sample 12) [7]; 304.205360 (iTRAQ 8-plex) for the mouse embryonic fibroblasts (sample 6) [3] and dendritic cells (sample 25) [7]; and 229.162932 (TMT 10-plex) for the labeled human T lymphocytes (sample 20), the human aortic intima and media layers (samples 10 and 11) [6] and the aortic tissue from Marfan syndrome patients (sample 9). The so-obtained peptide-spectrum matches (PSMs) and the differences between the observed precursor mass and the theoretical mass of the corresponding unmodified peptide (Δmass) were analyzed with the publicly available software package SHIFTS [12] version 0.3.1 (<https://github.com/CNIC-Proteomics/SHIFTS-4/releases/tag/‌v0.3.1>). In brief, SHIFTS comprises a set of software modules that i) sequentially recalibrate experimental mass and Δmass values using a high-quality PSM subset (0.15 minimum corrected Xcorr, 20 ppm tolerance); ii) model Δmass values into a histogram grouped by Δmass bins; iii) assign PSMs to Δmass peaks (defined by the frequency and slope of each bin); iv) calculate PSM global, local (1-Da binning) and peak false discovery rate (FDR); v) match Δmass values against a list of known mass shifts (most of which are taken from the Unimod database [13]); vi) check whether the Δmass could be explained by peptide truncation; and vii) check if Δmass values have been located to the right position in the PSM.

**References**

1.	Rojas-Gomez, A., et al., *Chaperonin CCT controls extracellular vesicle production and cell metabolism through kinesin dynamics.* J Extracell Vesicles, 2023. **12**(6): p. e12333.

2.	Martin-Cofreces, N.B., et al., *The chaperonin CCT controls T cell receptor-driven 3D configuration of centrioles.* Sci Adv, 2020. **6**(49).

3.	Bonzon-Kulichenko, E., et al., *Improved integrative analysis of the thiol redox proteome using filter-aided sample preparation.* J Proteomics, 2020. **214**: p. 103624.

4.	Almeida-Marques, C., et al., *Secretome processing for proteomics: A methods comparison.* Proteomics, 2024: p. e2300262.

5.	Sanchiz, A., et al., *The Experimental Proteome of Leishmania infantum Promastigote and Its Usefulness for Improving Gene Annotations.* Genes (Basel), 2020. **11**(9).

6.	Martinez-Lopez, D., et al., *Complement C5 Protein as a Marker of Subclinical Atherosclerosis.* J Am Coll Cardiol, 2020. **75**(16): p. 1926-1941.

7.	Izquierdo-Serrano, R., et al., *Extracellular vesicles from Listeria monocytogenes-infected dendritic cells alert the innate immune response.* Front Immunol, 2022. **13**: p. 946358.

8.	Torralba, D., et al., *Priming of dendritic cells by DNA-containing extracellular vesicles from activated T cells through antigen-driven contacts.* Nat Commun, 2018. **9**(1): p. 2658.

9.	Bache, N., et al., *A Novel LC System Embeds Analytes in Pre-formed Gradients for Rapid, Ultra-robust Proteomics.* Mol Cell Proteomics, 2018. **17**(11): p. 2284-2296.

10.	Hulstaert, N., et al., *ThermoRawFileParser: Modular, Scalable, and Cross-Platform RAW File Conversion.* J Proteome Res, 2020. **19**(1): p. 537-542.

11.	Laguillo-Gomez, A., et al., *ReCom: A semi-supervised approach to ultra-tolerant database search for improved identification of modified peptides.* J Proteomics, 2023. **287**: p. 104968.

12.	Bagwan, N., et al., *Comprehensive Quantification of the Modified Proteome Reveals Oxidative Heart Damage in Mitochondrial Heteroplasmy.* Cell Rep, 2018. **23**(12): p. 3685-3697 e4.

13.	Creasy, D.M. and J.S. Cottrell, *Unimod: Protein modifications for mass spectrometry.* Proteomics, 2004. **4**(6): p. 1534-6.
